# *Vibrio cholerae* may be transmitted to humans from bullfrog through food or water

**DOI:** 10.1101/2021.04.09.439145

**Authors:** Yibin Yang, Xia Zhu, Yuhua Chen, Yongtao Liu, Yi Song, Xiaohui Ai

**Affiliations:** Yangtze River Fisheries Research Institute, Chinese Academy of Fishery Sciences, Wuhan 430223, China; The Key Laboratory for quality and safety control of aquatic products, Ministry of Agriculture, Beijing 100037, China; Shanghai Ocean University, Shanghai 201306, China; Department of Gastroenterology, Zhongnan Hospital of Wuhan University, Wuhan 430227, China; Hubei Clinical Center & Key Lab of Intestinal & Colorectal Diseases, Wuhan 430227, China

**Keywords:** *Vibrio cholerae*, Bullfrog, Food safety, Fatal threat, Route of transmission

## Abstract

Bullfrog is one of the most important economic aquatic animals in China. It is widely cultured in southern China, and is a key breed recommended as an industry of poverty alleviation in China. During recent years, a fatal bacterial disease has often been found in cultured bullfrogs. The clinical manifestations of the diseased bullfrogs were severe intestinal inflammation and even anal prolapse. A bacterial pathogen was isolated from the diseased bullfrog intestines. The bacterium was identified as *Vibrio cholerae* using morphological, biochemical and 16S rRNA phylogenetic analysis. In this study, *V. cholerae* was isolated and identified from diseased bullfrogs for the first time, providing a basis for the diagnosis and control of the disease. At the same time, it was also found that *V. cholerae* may be transmitted to humans from bullfrogs through bullfrog food and aquaculture water, creating a serious threat for human health. Therefore, society should pay attention to the modes of transmission of *Vibrio cholerae* from bullfrog and formulate reasonable safety measures to avoid disasters.

## 1. Introduction

Bullfrog belongs to the Ranae and Rana, family and genus, respectively, of the order Anura and class Amphibia of the phylum Chordata. It is the most popular large edible frog in the world. It is named based on the loud sound and its resemblance to the noise made by cattle (Gao, 2016). The original distribution of the Bullfrog is to the east of the Rocky Mountains in the United States, 30° to 40° north latitude area and southern Ontario and Quebec in Canada. It is the largest frog found in North America (Nori, Urbina-Cardona, Loyola, Lescano & Leynaud, 2011). The adult frog is generally 8-12 cm in length and can reach a maximum weight of 2 kg. Although bullfrogs are about 300 million years old, its artificial culture spans only about a hundred years. Due to its delicious taste, high protein content, low fat and low cholesterol content, its skin, oil, hormone, gland and bile are of economic value, and are used as important raw materials in aquaculture, medicine and other industries (Gao, 2016). Therefore, bullfrog has been favored by many consumers since it was first introduced into China from Cuba in 1959 (Ding et al., 2020). It is one of the main economically valuable aquaculture animals in China. At present, bullfrog breeding is mainly distributed in Guangdong, Fujian, Zhejiang, Jiangxi, Hainan, Anhui, Jiangsu, Hunan, Hubei, Sichuan and other southern regions (Gao, 2016).

During the raring process of bullfrogs, various techniques are used by farmers to improve yield, and little attention has been paid to the carrying capacity of the waterbodies, increasing the seriousness of diseases year by year along with the increase in breeding density. Along with the continuous increase of the scale and density of bullfrog breeding, many problems, such as the shortage of biologically healthy food, the degradation of germplasm resources, the deterioration of the breeding environment, and a lack of breeding technology, have emerged. The diseases encountered during bullfrog breeding are on the rise and are becoming increasingly more detrimental, with large-scale outbreaks occurring from time to time, which has seriously hindered the development of the industry. At present, the main pathogens of bullfrog diseases are bacteria, viruses, and parasites (Ding et al., 2020; Pu et al., 2019; Yu et al., 2013). Due to the characteristics of many types of pathogens, complex and diverse causes, rapid spread and high mortality, bacterial diseases are the most harmful to the bullfrog breeding industry, and have become the focus of prevention and control during the process of bullfrog breeding (Yu et al., 2013).

During recent years, a strange disease has often broken out in bullfrog farms in the Zhangzhou area of Fujian Province, which is commonly known as anorectal disease by local farmers. The main clinical symptoms of diseased bullfrogs are anal abscission, rotten feces, signs of severe dyspepsia, which has been confirmed as severe enteritis through diagnosis. To find out the cause of the disease in bullfrogs as soon as possible, and to formulate prevention and control measures, the etiology of the disease was studied in bullfrogs with typical symptoms. Through bacteriological studies, several dominant strains were isolated from different batches of samples. Further studies found that the dominant bacteria in most samples was the same based on morphology and dominance. According to Koch criterion, the pathogenicity of the bacteria isolated bullfrog was studied. The results showed that the isolated strain was a pathogen of bullfrog that could also cause fatal diseases in humans. In history, the isolates have caused epidemics that have resulted in hundreds of millions of human deaths. The results of this study provide important methods for the diagnosis and control of this emerging disease in bullfrog and warn that there may be fatal risks associated with the human consumption of bullfrog products or contact with water used for breeding bullfrog.

## 2. Materials and methods

### 2.1 Sampling

Bullfrogs with typical symptoms were collected multiple times from Zhangzhou, Fujian Province. The diseased bullfrogs with typical symptoms were put in net bags and brought back to the laboratory for diagnosis and pathogen isolation. Bullfrogs (50 ± 2 g) were purchased for infection experiments from farms without a history of disease. The purchased bullfrogs showed a strong jumping ability with no scars on their bodies. The healthy bullfrogs were kept in buckets for 7 days, and the water used for breeding was not higher than the neck of the frogs. Seven days later, bullfrogs were used in infection tests if they seemed to be normal.

### 2.2 Pathogen isolation

A light microscope was used to observe the intestinal tract to identify typical symptoms of parasitic or fungi infections. Bacterial isolation was performed in a secondary biosafety cabinet (ESCO, Singapore). The anesthetized bullfrogs were placed on ice and disinfected with 75% ethanol before dissection. After the intestinal and visceral tissues of each bullfrog were allowed contact with the inoculation ring, the inoculation ring was crossed on the agar plate of the brain heart extract (BHI; Difco, USA), and the plate was cultured at 28°C for 24 h. The dominant strain was selected and then purified. The purified strain was temporarily named NW01. The purified strain was mixed with 15% glycerol and frozen at −80°C as a standby.

### 2.3 Biochemical characterization of bacterial isolates

The isolate NW01 was inoculated on agar medium plates with brain heart extract and cultured at 28°C for 24 hours. The isolated strains were stained using gram, and the physicochemical indexes of the isolated strains were determined through micro biochemical identification based on the manual for the identification of common bacterial systems (Dong & Cai, 2001).

### 2.4 16S ribosomal RNA sequencing analysis of the isolates

The isolated strains were inoculated into agar medium plates with brain heart extract and cultured at 28°C for 18 h. A single colony was selected and placed in 10 μL of sterile water, and then it was blown evenly to be used as a template for PCR.

The universal primers used for 16S rRNA were 27F: 5’- agagtttgatc (c/a) tggctcag-3’, and 1492R: 5 ‘- ggttaccttgttacgatt-3’ (Polz & Cavanaugh, 1998). PCR reaction system: 50 μL of 2 × Taq PCR mix, 47 μL of ddH2O, 1 μL of upstream and downstream primers, and 1 μL of the template. Reaction conditions: 35 cycles of denaturation at 95°C for 1 min, equilibrium at 98°C for 15 s, annealing at 55°C for 30 s, extension at 72°C for 2 min, followed by incubation at 72 °C for 10 min. The amplified product was verified using 1% agarose gel electrophoresis as the target fragment size and then sent to Shanghai bioengineering for purification and sequencing. The 16S rRNA gene sequence of the NW01 strain was added into NCBI for comparison. The 16S rRNA sequences of *Vibrio* and important aquatic pathogens were selected and analyzed using cluster x software. The phylogenetic tree was constructed using the neighbor joining method using mega 6.0 software and the confidence interval of the bootstrapping was 10,000 times.

### 2.5 Pathogenicity

According to Koch’s rule, the regression infection experiment was designed to determine whether the isolate was pathogenic to bullfrogs, to confirm whether the isolate was the pathogen that caused disease in bullfrogs. The isolated strains were cultured in brain heart extract medium at 28°C for 18 h. The bacteria (sigma, 3k15) were collected at 4000 rpm for 5 minutes at 4°C, and then the bacterial mass was resuspended in a sterile PBS buffer. The concentration of the bacterial suspension was adjusted to about 10^4^, 10^6^ and 10^8^ CFU/mL. One hundred and twenty healthy bullfrogs were randomly divided into four groups (A, B, C and D) with 30 in each group. Group A, B and C were used as the experimental groups, while group D was used as the control group. The bullfrogs in group A-C were intraperitoneally injected with 0.1 mL of the bacterial suspension, and the concentrations of the bacterial suspensions used were 10^8^, 10^6^ and 10^4^ CFU/mL respectively, that is that the injection doses were 10^7^, 10^5^ and 10^3^ CFU/frog, while the bullfrogs in group D were injected with the same dose of PBS at the same site. During the experiment, the air temperature was controlled at 24-26°C and ventilation was kept constant. Fully aerated tap water was used for breeding. The water used for breeding did not exceed the neck of the bullfrogs. Each bucket was covered with a gray cover to prevent the frog from escaping. The water used for breeding was changed every day. The state of the experimental bullfrogs were observed until death. Clinical symptoms and mortality were recorded every day, and bacteria were isolated from the dying bullfrog and purified. The purified bacteria were identified using 16S rRNA sequencing.

### 2.6 Analysis of drug sensitivity of the NW01 strain

A standard NCCLS antimicrobial susceptibility test was conducted using the paper diffusion method to analyze the antimicrobial susceptibility of the isolates (CLSI, 2006). The isolate NW01 was inoculated into a nutrient broth and cultured at 28°C for 24 hours at 200 rpm. The bacterial suspension was diluted with PBS to a concentration of 10^7^ CFU/mL. 100 μL of the bacteria suspension was used to coat MH agar, and the selected drug sensitive paper was pasted on the plate. The drug content of the paper is shown in the table. The plate was incubated at 28°C for 24 h, and the size of the inhibition zone was measured.

### 2.7 Serotype identification of the isolate

The serotype of the isolated bacteria was identified based on the method used for the *Vibrio cholerae* O antigen diagnostic serum. First, the suspension of bacteria to be tested was dripped onto a clean slide, and then a single drop of *V. cholera* O1 group, O1 group Oryza type, O1 group Ogawa type, O139 group diagnostic serum was dripped onto the suspension of bacteria to be tested, and mixed evenly, and observed 1 min later. Meanwhile, physiological saline was used as the control. Agglutination at 2 + or more was considered as a positive result.

## 3. Results

### 3.1 The epidemic time and clinical symptoms of diseased bullfrog

Epidemiological investigations showed that the disease affected the entire breeding cycle of the bullfrogs, especially during the high temperature season. The increase in feeding intensity (Fig. 1A-C) made the intestinal tract of bullfrogs more prone to inflammation, leading to anal abscission (Fig. 2). The diseased bullfrogs showed no obvious symptoms on the surface of their body. Dissection showed that the diseased bullfrogs suffered slight congestion and swelling in its viscera, severe intestinal inflammation, and rotten feces. Based on the symptoms, the disease was name as bullfrog enteritis. The weight of the diseased bullfrogs ranged from 100 to 1000 g, and there were no significant individual differences. During the investigation, the incidence rate of enteritis in bullfrogs was high, but the mortality rate was not high, indicating a chronic disease. Enteritis can lead to indigestion, malnutrition, and intestinal mucosa damage of the bullfrog, which makes it easier for other pathogens, such as *Streptococcus* infection, to occur. In addition, enteritis can lead to anal prolapse of bullfrogs, seriously affecting the appearance of commercial bullfrogs, resulting in exceptionally large economic losses for farmers. No parasite or fungus infection was found in bullfrogs as observed under a light microscope. After the bacteriological study, a strain of bacteria was isolated and purified, and was temporarily named NW01.

**Figure 1.**
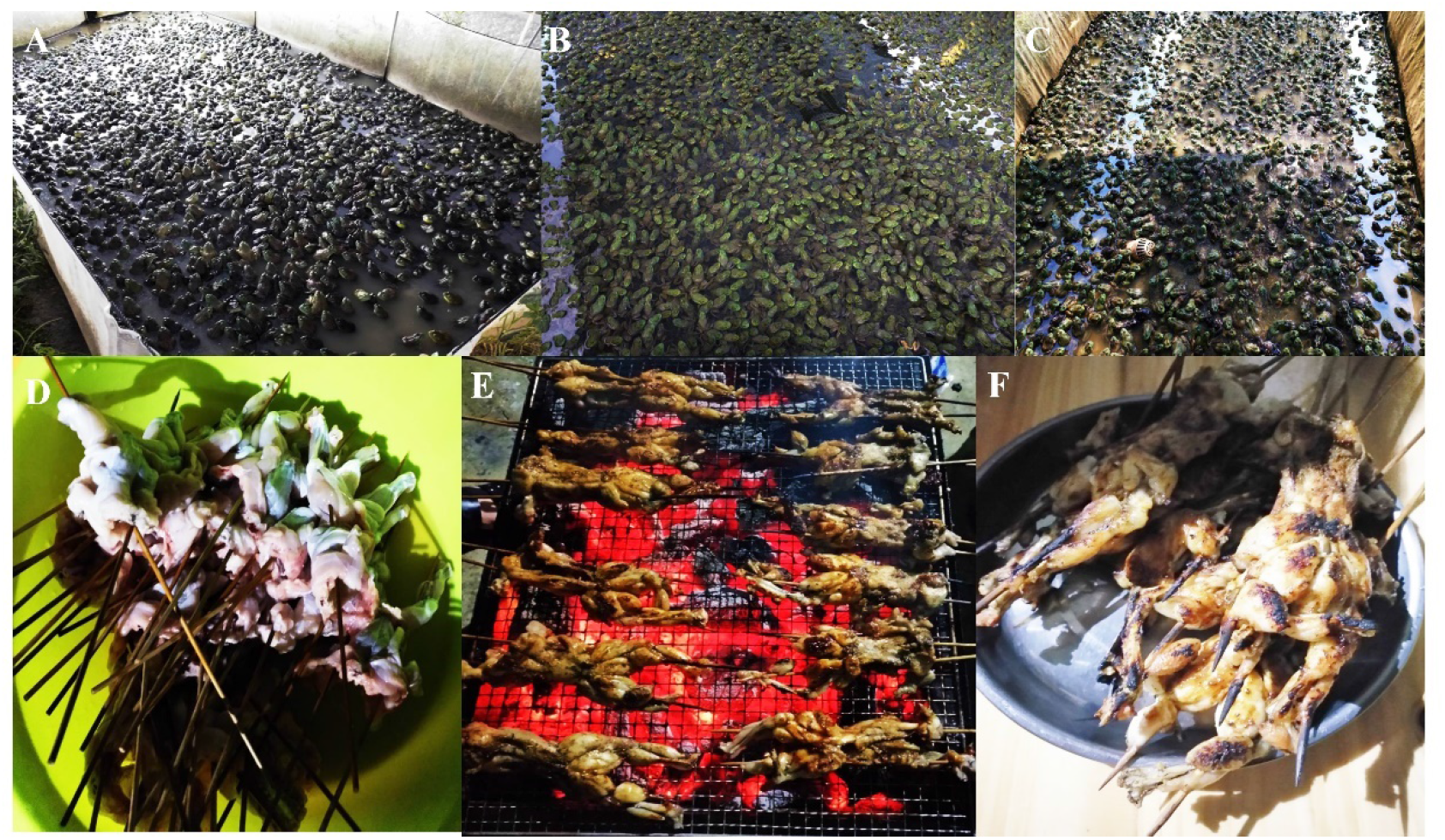
Cultivation and feed for bullfrog: A-C: cultivation; D-E: feed

**Figure. 2.**
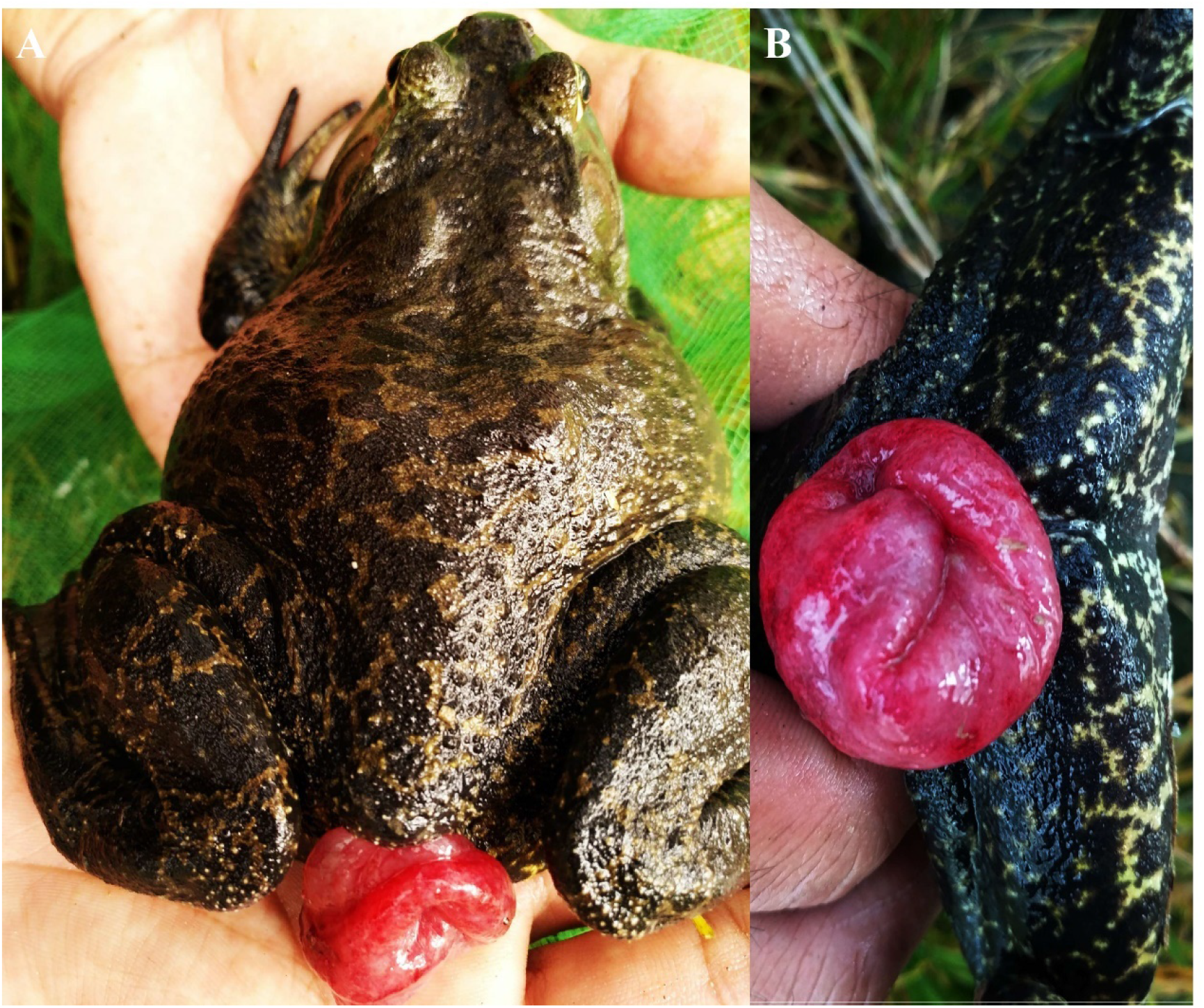
A diseased bullfrog

### 3.2 Biochemical characterization and molecular identification of the bacteria

The result of Gram staining showed that the isolate NW01 was red, indicating that it was gram negative. The specific physical and chemical characteristics of the bacteria are shown in Table 1. The physical and chemical characteristics showed that NW01 was *Vibrio cholerae*. The 16S rRNA gene of NW01 was amplified using universal primers, and the 16S rRNA fragment of NW01 was about 1500 bp long, which is in line with the expected size. The 16S rRNA gene sequences (GenBank accession number: MT126343) of the isolated strains were added into a gene library and analyzed using the ncbi-blast program. The results showed that the isolates had the highest homology with *V. cholerae*. The 16S rRNA gene sequences of several *Vibrio* species and important aquatic pathogens were selected to construct a phylogenetic tree based on the 16S rRNA gene sequences, as shown in figure 3. The results showed that the isolates and *V. cholerae* were clustered into one branch. Therefore, the combination of physical and chemical characteristics and gene analysis of the isolate confirmed that NW01 was *V. cholerae*.

**Figure 3.**
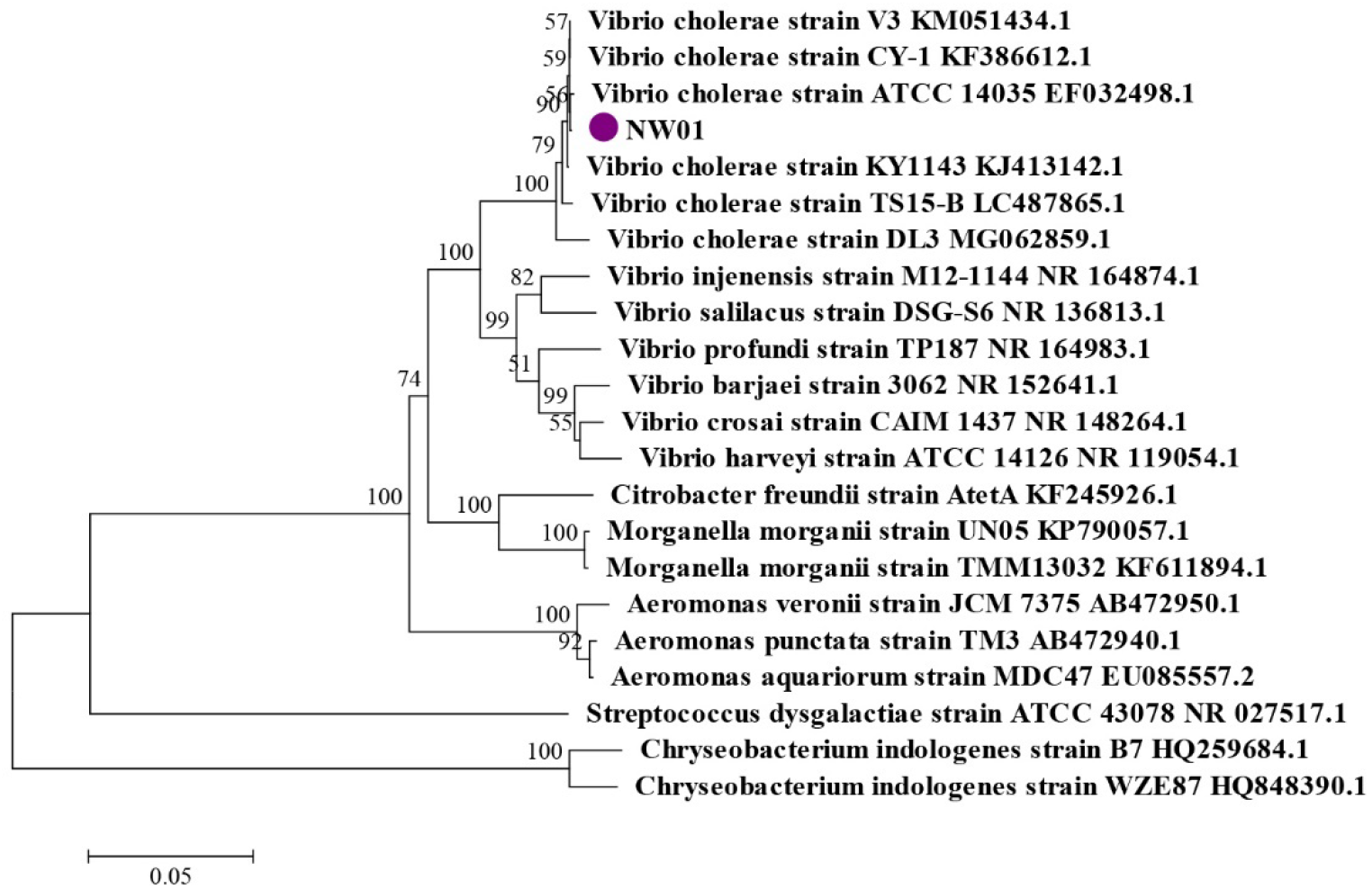
The phylogenetic tree for the 16s rRNA sequence of NW01

**Table 1.**
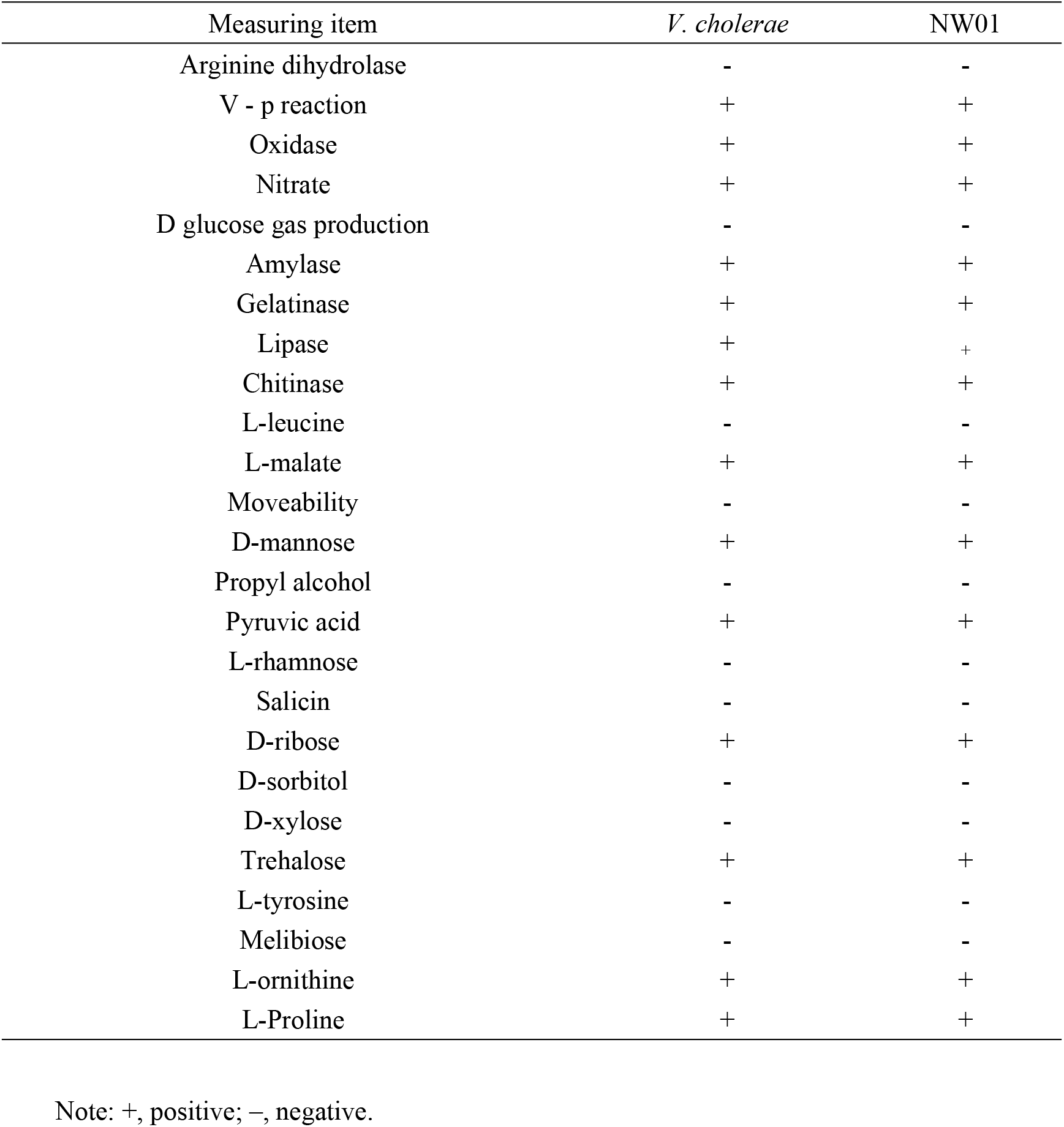
Physiological and biochemical characteristics of the NW01 strain

| Measuring item | *V. cholerae* | NW01 |
| --- | --- | --- |
| Arginine dihydrolase | - | - |
| V - p reaction | + | + |
| Oxidase | + | + |
| Nitrate | + | + |
| D glucose gas production | - | - |
| Amylase | + | + |
| Gelatinase | + | + |
| Lipase | + | + |
| Chitinase | + | + |
| L-leucine | - | - |
| L-malate | + | + |
| Moveability | - | - |
| D-mannose | + | + |
| Propyl alcohol | - | - |
| Pyruvic acid | + | + |
| L-rhamnose | - | - |
| Salicin | - | - |
| D-ribose | + | + |
| D-sorbitol | - | - |
| D-xylose | - | - |
| Trehalose | + | + |
| L-tyrosine | - | - |
| Melibiose | - | - |
| L-ornithine | + | + |
| L-Proline | + | + |

Note: +, positive; -, negative.

### 3.3 Pathogenicity

In the pathogenicity study of the isolate, the experimental groups all died to varying degrees (Fig. 4). The mortality rates of group A, B and C were 100%, 80% and 23.33%, respectively. The dead bullfrogs showed similar symptoms to natural disease, while bullfrogs in the control group did not get sick or die. *V. cholerae* was isolated again from the dying bullfrogs. The infection experiment was performed in accordance with Koch’s law, and the results indicated that *V. cholerae* was the pathogen that had caused bullfrog enteritis.

**Figure 4.**
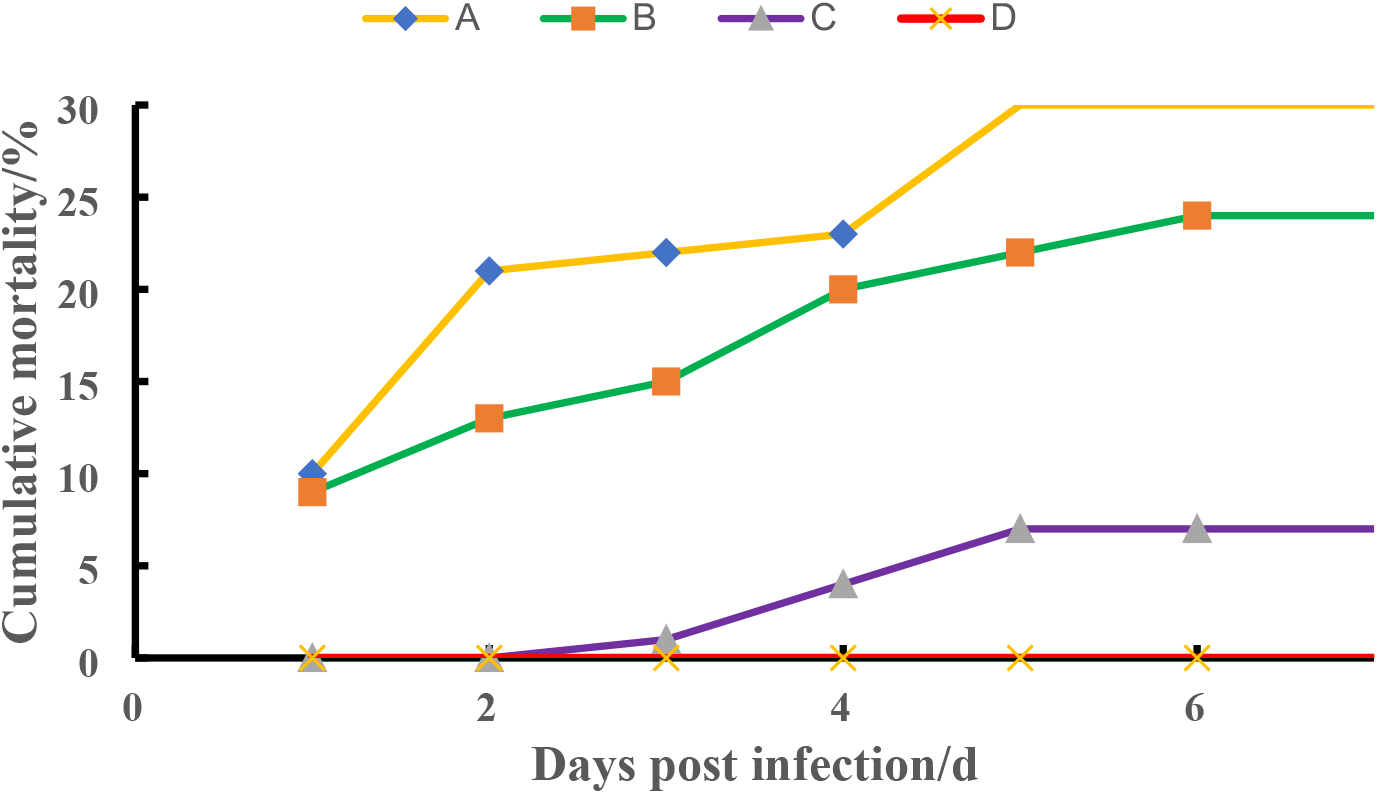
The pathogenicity of the healthy bullfrogs experimentally infected with 10^7^ (A), 10^5^ (B), 10^3^ (C) CFU/bullfrog doses of the isolated NW01 strain or 0.1 ml of PBS (D).

### 3.4 Drug sensitivity tests for the NW01 strain

The sensitivity of NW01 to 20 antibiotics was determined. The results showed that NW01 was resistant to β - lactams, aminoglycosides, macrolides, tetracyclines and sulfonamides, but sensitive to cephalosporins, quinolones and amido alcohols. Among the sensitive drugs, neomycin, doxycycline and florfenicol are allowed to be used in aquaculture (Table 2). Therefore, neomycin can be used for a course of 7 days to control the spread of the disease, but it cannot easily improve bullfrog anal prolapse. Therefore, neomycin was selected to treat bullfrog enteritis. A

**Table 2.**
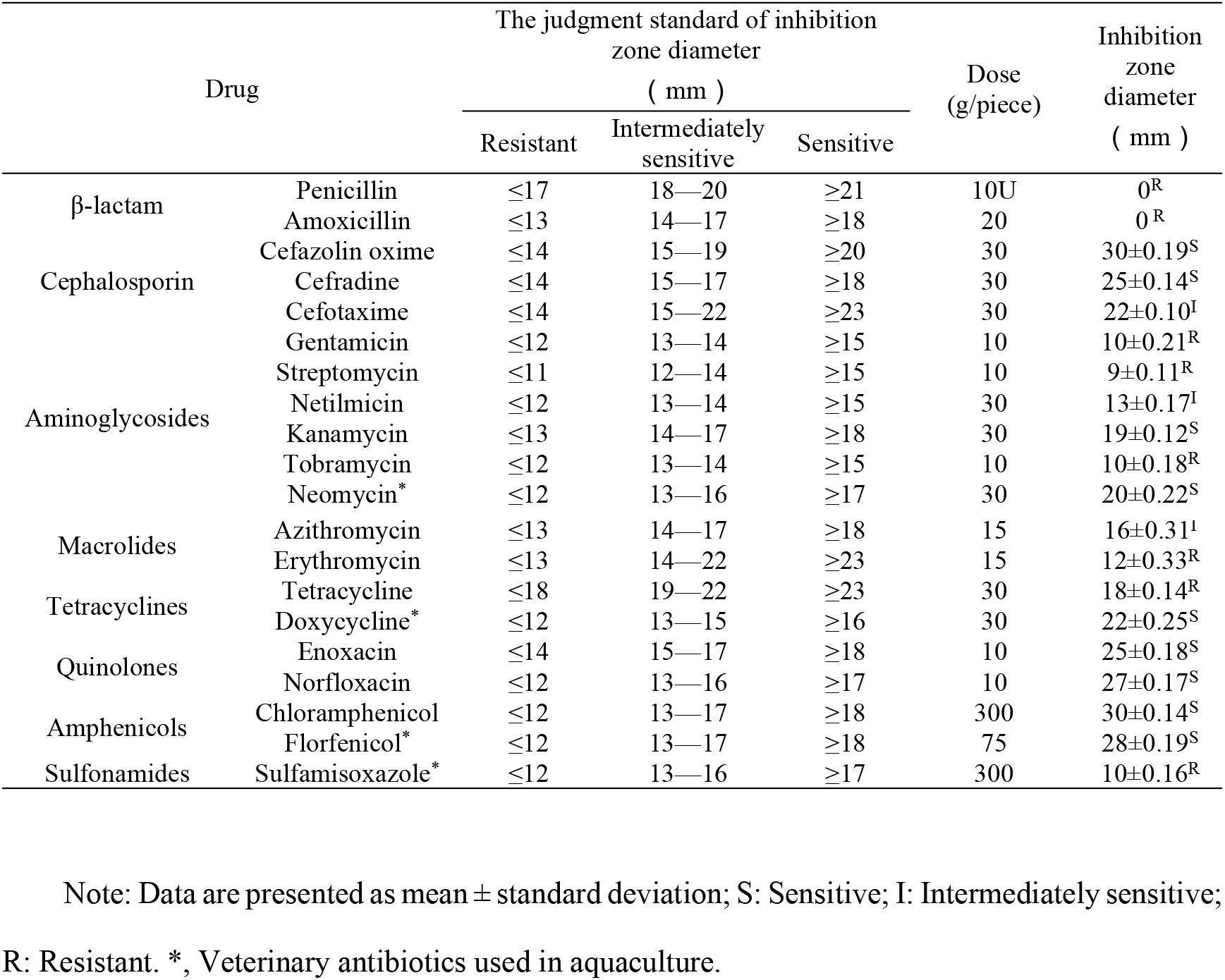
Susceptibility of NW01 to antibiotics

### 3.5 Serotype identification of the isolated bacteria

The O antigen slide agglutination test was used to determine that the isolated strain NW01 did not agglutinate with the *V. cholerae* O1 group, O1 group rice leaf type, O1 group Ogawa type, or O139 diagnostic serum, and the isolate was further identified as non-O1/non-O139 group *V. cholerae*.

## 4 Discussion

The breeding of frogs has a long history in China. Frogs are bred not only as food, but also to harvest a variety of industrial raw materials and provides good economic benefits. Bullfrog is an important representative species (Zhang, 2015). Along with large developments in the bullfrog breeding industry, diseases have began to occur more frequently during the process of breeding. Due to largescale domestic bullfrog breeding, the reports of bullfrog diseases have been mainly concentrated in China (Pu et al., 2019; Han et al., 2016), Overseas, only South Korea, France, North America and a few other countries have reported of the same. The main pathogens of bullfrog diseases include viruses, bacteria, and parasites (Candido et al., 2019; Jaÿ et al., 2020; Khalifa & Bekhet, 2018; Kim, Koo, Park, Kwon, Park, & Ecology, 2016; Landsberg, Kiryu, Tabuchi, Waltzek, & Pessier, 2013; Oliveira, Alfaia, Ikari, Tavares, & Ferreira, 2019). The frequent occurrence of diseases in bullfrog has caused great economic losses, and diseases are becoming a major bottleneck in the development of the bullfrog industry.

In this study, the dominant strain, NW01, was isolated from diseased bullfrogs. The isolate NW01, was identified as *V. cholerae* through biochemical identification, 16S rRNA sequence analysis and construction of a phylogenetic tree. The regression infection experiment confirmed that *V. cholerae* was the pathogen that caused bullfrog enteritis. The results showed that *V. cholerae*, a zoonotic bacterium, caused the first infection in the bullfrogs, which led to a great epidemic of diseases and caused great economic losses.

Based on the sensitivity of the isolates to 20 types of antibiotics, neomycin was selected to be used for clinical prevention and control in this study, and the spread of the epidemic was controlled in time. However, it cannot induce a good therapeutic effect on bullfrog anal prolapse, as it is a serious condition that is difficult to improve.

*V. cholerae* belongs to *Vibrio* family and can be divided into 139 serogroups. Among them, O1 and O139 can cause cholera in humans. O1 group and O139 group can cause cholera mainly because they carry the cholera toxin (Faruque, Albert, & Mekalanos, 1998; Sánchez & Holmgren, 2011), which can activate adenylate cyclase in intestinal epithelial cells, resulting in the secretion of Cl^-^ ions and impairment of Na^+^ ion absorption. Water enters the intestinal cavity with ions, causing severe watery diarrhea, which leads to human death. Cholera is an ancient and widespread infectious disease that mainly manifests as severe vomiting, diarrhea, water loss, high mortality, and is an international quarantine class A infectious disease. Since 1817, there have been seven cholera pandemics worldwide, causing hundreds of millions of human deaths. However, non-O1 and non-O139 *V. cholerae* may carry other virulence factors, which are widely distributed in the water environment. They can usually cause human gastrointestinal inflammation and may sometimes cause extraintestinal infections, such as meningitis, sepsis, and wound infections (Hounmanou et al., 2016).

*V. cholerae* widely exists in all types of waterbodies (Daboul, Weghorst, Deangelis, Plecha, & Matson, 2020; Hounmanou et al., 2016). It has been reported that *V. cholerae* can infect aquatic animals. At present, it has been reported that *V. cholerae* can infect fish (Reddacliff, Hornitzky, Carson, Petersen, & Zelski, 2010; Rehulka, Petras, Marejkova, & Aldova, 2015), shrimps (Li et al., 2019; Zhou et al., 2020), and other aquaculture animals (López-Hernández et al., 2015; Kawai, Ota, Takemura, Nakai, & Maruyama, 2020). The cause of cholera epidemics are extraordinarily complex, and it is unclear how it spreads, while the reason for seasonal epidemic peaks in epidemic areas are also unknown. However, it is an indisputable fact that cholera is caused by *V. cholerae* (Byun, Jung, Chen, Larios Valencia, & Zhu, 2020).

Since the transmission mechanism of *V. cholerae* is not clear, it has been thought that aquatic animals are infected with non-O1 and non-O139 *V. cholerae*. The *V. cholerae* isolated from bullfrogs were also of the non-O1 group and non-O139 group, which can cause symptoms of intestinal inflammation and even anal prolapse, results in low mortality and a long duration of survival, but a high incidence rate.

However, further research has shown that the O1 group and O139 group of *V. cholerae* have been reported in aquatic animals, such as reports of O1 group found in tilapia (Hounmanou et al., 2019) and O139 group found in loach and shrimp (Joseph, Murugadas, Reghunathan, Shaheer, Akhilnath, & Lalitha, 2015; Chen et al., 2016). These reports indicate that *V. cholerae* in aquatic animals is not all non-O1 group and non-O139 group, but may also be O1 group and O139 group, which can cause cholera outbreaks. Therefore, it is obvious that *V. cholerae* is an important zoonotic bacterium. Although there are no reports of human cholera outbreaks caused by O1 and O139 from aquatic animals, it is unknown whether such cholera outbreaks will occur in the future. As an open community similar to that of wild animals, aquatic animals are likely to serve as the resource repository for zoonotic bacteria, such as *V. cholerae*. Further attention needs to be paid to determine whether the pathogen can spread, mutate and evolve among aquatic animals.

According to the author’s unpublished paper, *V. cholerae* from aquatic animals may spread through birds, aquatic products (food) and aquaculture water. At present, there are many methods in which bullfrogs are used as food, among which hot pot bullfrogs and barbecued bullfrogs are the two main ways, and the bullfrogs prepared using these methods may be eaten without being fully cooked (Fig. 1D-F). In addition, there are health risks and cross infection risks in Chinese restaurants. At present, on one hand, there is no systematic policy for the detection of pathogenic microorganisms in aquatic products in China, and there are also some loopholes. On the other hand, to reduce economic losses, sick bullfrogs are sold at a lower price. These results indicate that bullfrogs are likely to be infected with *V. cholerae* through aquatic products. Other studies have found *Salmonella* and microsporidia in bullfrogs, which are also serious zoonotic pathogens (Ding et al., 2020; Zhang et al., 2015). Therefore this series of findings is worthy of attention.

Bullfrogs belongs are amphibian but water should not cover its neck during the process of breeding, therefore bullfrog pools contain only a small amount of water. Bullfrogs eat a lot and discharge a lot of feces and urine. Therefore, the water used for breeding bullfrogs is often black, with a foul smell. The ammonia nitrogen index is often dozens of times higher than that which should be used for breeding, and is full of organic matter. These observations show that the aquaculture water and living environment of bullfrog provides rich nutrition for the reproduction of *V. cholerae*. Therefore, the water used in bullfrog breeding is likely to act as a culture medium of *V. cholerae* and becomes the mother liquor of *V. cholerae*. The mother liquor is directly discharged into natural waterbodies without treatment and can permeate drinking water sources, resulting in severely detrimental consequences.

Although only non-O1 and non-O139 *V. cholerae* were found in bullfrogs in this study, it is possible that *V. cholerae* can become O1 and O139 serotypes through gene transfer under the current open culture practice of bullfrogs (Bai, Ke, Consuegra, Liu, & Li, 2012), with further enhanced risk. Bullfrog is also a biologically invasive species in China and has a strong survival ability in the wild, resulting in the further spread and variation of the pathogen. Chinese people have always been fond of eating wild animals, which also provides an important method for the spread of pathogens. Therefore, the author calls for the strengthening of the monitoring of *V. cholerae* and other zoonotic pathogens in edible bullfrogs to ensure the quality and safety of bullfrog aquatic products. At the same time, bullfrog breeding wastewater should be treated to a great extent to avoid *V. cholerae* pollution of natural waterways, which will lead to human health concerns and fatalities.

## ACKNOWLEDGEMENTS

This work was supported by the China Agriculture Research System CARS-46 and Modern Agricultural Talent Support Plan (2016-139). Thanks are also due to the anonymous reviewers who provided detailed comments that helped to improve the manuscript.

## ETHICAL APPROVAL

The authors confirm that the ethical policies of the journal, as noted on the journal’s author guidelines page, have been adhered to. No ethical approval was required as human or animal subjects were not involved in this study.

## DATA AVAILABILITYS TATEMENT

All data generated or used during the study appear in the submitted article.

